# Tradeoffs for a viral mutant with enhanced replication speed

**DOI:** 10.1101/2021.03.24.436823

**Authors:** Matthew R. Lanahan, Julie K. Pfeiffer

**Affiliations:** Department of Microbiology, University of Texas Southwestern Medical Center, Dallas, Texas, USA

**Keywords:** coxsackievirus, replication speed, fitness, capsid

## Abstract

RNA viruses exist as genetically heterogeneous populations due to high mutation rates and many of these mutations reduce fitness and/or replication speed. However, it is unknown whether mutations can increase replication speed of a virus already well adapted to replication in cultured cells. By sequentially passaging coxsackievirus B3 in cultured cells and collecting the very earliest progeny, we selected for increased replication speed. We found that a single mutation in a viral capsid protein, VP1-F106L, was sufficient for the fast-replication phenotype. Characterization of this mutant revealed quicker genome release during entry compared to wild-type virus, highlighting a previously unappreciated infection barrier. However, this mutation also reduced capsid stability *in vitro* and reduced replication and pathogenesis in mice. These results reveal a tradeoff between overall replication speed and fitness. Importantly, this approach— selecting for the earliest viral progeny—could be applied to a variety of viral systems and has the potential to reveal unanticipated inefficiencies in viral replication cycles.

**Significance:** Viruses have characteristic replication speeds within a given cell type. Many factors can slow the rate of viral replication, including attenuating mutations and host antiviral responses. However, it has been unclear whether it would be possible to “speed up” a virus that already replicates efficiently in a specific cell type. Here, we selected for a mutant coxsackievirus with enhanced replication speed by sequentially harvesting the very earliest progeny in multiple rounds of selection. A single mutation conferred the fast replication phenotype. While this mutant virus has enhanced replication in cultured cells due to faster genome uncoating, it was attenuated in mice. These results highlight selective pressures that shape viral populations in different environments.

## Introduction

The growth rate of an organism is a selectable trait that depends on both intrinsic and extrinsic factors. Humans have selected for organisms with faster growth rates, including chickens, hogs, maize, and soybeans [1]. Similar practices have been used to select for bacteria, fungi, and viruses that replicate efficiently in laboratory conditions [2]. Determining the proper extrinsic factors--such as growth media, pH, temperature— that facilitate efficient replication was foundational for our ability to cultivate microorganisms. In the past several decades, many viruses have been passaged in cultured cells to generate lab-adapted viruses that replicate efficiently in a given cell type [3-7]. More recently, several groups have uncovered genetic changes in bacteria [8, 9] and fungi [10, 11] that confer faster replication. These mutants are typically discovered through forward genetic selection by serial passaging or random mutagenesis.

Genetic variation is a key component of natural selection and evolution. RNA viruses have high mutation rates, and it is thought that this is a highly selected for and evolved trait [12, 13]. Many RNA viruses make about one error per 10,000 nucleotides due to a lack of proofreading by the RNA-dependent RNA polymerase [13]. As a result, RNA virus populations are highly diverse. Most mutations are deleterious [14]. Therefore, viruses with increased mutation frequencies typically lose fitness due to “error catastrophe” [15, 16]. RNA viruses with lower-than-normal mutation frequencies also face a fitness cost [17-19], but there are differing hypotheses as to why this occurs. One hypothesis is that a lower mutation frequency is biochemically costly for the viral polymerase and reduces overall replication speed, which in turn, reduces fitness [20, 21]. Another hypothesis is that genetic diversity enhances fitness through increased adaptation in the presence of selective pressure. This idea is supported by studies in which anti-mutator strains are attenuated *in vivo* [13-15], but attenuation can be overcome by increasing mutation frequency [19]. Taken together, these findings suggest that viruses have evolved an optimized mutation rate that is dependent on many factors including viral adaptability, population fitness, and replication speed.

Coxsackievirus B3 (CVB3) is a useful model system to study how selective pressures influence viral populations. CVB3 is non-enveloped RNA virus in the *Picornaviridae* family. CVB3 can cause a wide range of pathologies, including myocarditis, type I diabetes, and meningitis, but most infections are mild [22]. CVB3 has a relatively fast replication cycle of 6-8 hours and replicates efficiently in both cell culture and mice. For polarized epithelial cells, CVB3 binds to Decay Accelerating Factor (DAF) on the cells surface, followed by translocation to CAR at tight junctions [23, 24]. In non-polarized cells, binding to DAF is not required since CAR is present on the cell surface [25]. Some CVB3 strains use heparin sulfate to facilitate attachment [26, 27]. Years ago, CVB3 strains were lab adapted such that they replicate efficiently in cultured cell lines such as HeLa, and infectious clones were generated from these cell culture adapted strains. One such strain, CVB3-H3, replicates very efficiently in HeLa cells, yielding a 1000-fold increase in viral titer over a single 8 hour cycle of replication [15]. While many CVB3 mutants have been characterized, nearly all are neutral or reduce viral fitness. While mutation acquisition can increase replication speed of attenuated/slow replicating picornaviruses [21], it is unclear if increasing replication speed of a “WT” virus is possible.

In this study, we used a forward genetic approach to select for CVB3 variants with faster replication speed. We found that a single mutation, F106L in the VP1 capsid protein, was sufficient for a fast-replication phenotype in HeLa cells. Characterization of the VP1-F106L mutant revealed that it had enhanced RNA release into cells during entry, which correlated with reduced virion stability. Although the VP1-F106L mutant had a fitness advantage over WT in cell culture, it was attenuated in mice. These studies reveal that it is possible to enhance replication speed of a virus, but this trait is likely selected against in nature to due to fitness costs in the presence of selective pressures within a host.

## Results

### Selecting for fast replicating CVB3 mutants

To determine whether a virus that replicates efficiently in cultured cells could evolve to replicate even faster, we performed serial passages of CVB3-H3 in HeLa cells and harvested the earliest progeny to enrich for potential “fast mutants”. Peak viral yield from single cycle replication of CVB3 in HeLa cells occurs at ∼8 hours post-infection. We infected HeLa cells at an MOI of 0.5 with a low-passage stock of CVB3 and collected cells at 4 hours post-infection to harvest the very earliest progeny, or 8 hours post infection to harvest progeny from a complete cycle of replication. Samples were then freeze-thawed to release intracellular virus, and half of the sample was used to initiate a new passage. This process was repeated, in triplicate, for 10 passages. After 10 passages, we performed a single cycle growth curve at an MOI of 0.1 to determine replication kinetics of the passaged samples and the input virus. Samples collected at 4 hours post-infection (4h P10) produced higher yields than samples collected at 8 hours post-infection (8h P10) or input stock virus (WT P0) at early time points in the replication cycle (Figure 1A). The largest titer increases for the 4h P10 samples were at early time points, 4h and 5h post infection, where 4h P10 samples had titers more than 10-fold higher than the stock/inoculum virus. These data suggest that the 4h P10 population had faster replication kinetics compared with WT P0 or 8h P10 populations.

**FIG 1.**
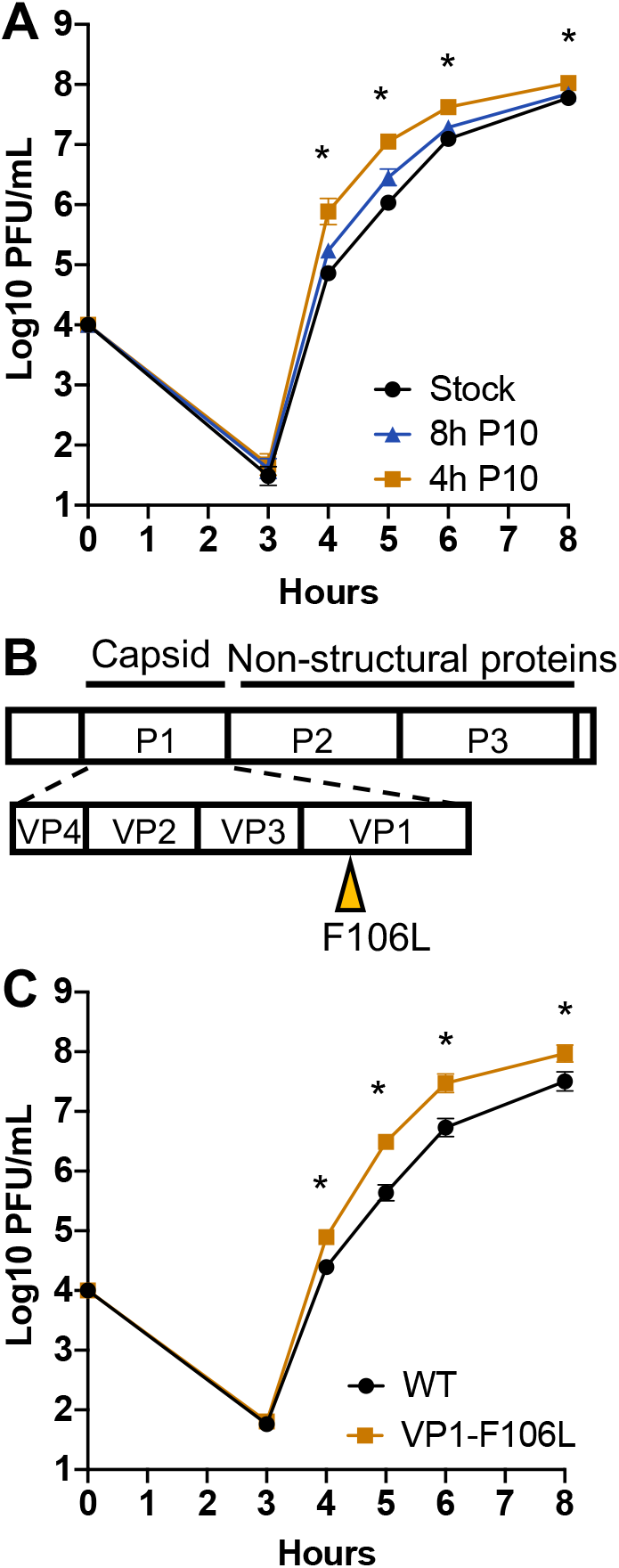
Selection for fast-replicating CVB3. (A) Single cycle growth curves for the initial stock virus, and viruses passaged 10 times for 4h or 8h (4h-P10 and 8h-P10). Viruses were used to infect HeLa cells at MOI=0.5, and progeny were quantified by plaque assay. Data are mean ± SEM (n≥11, ≥3 independent experiments). * p<0.05, two-way ANOVA (comparing 4h P10 and stock). (B) Consensus sequencing revealed that VP1-F106L was the only non-silent mutation found in the 4h P10 samples and not the 8h P10 samples. VP1-F106L was cloned into the infectious clone and used to generate VP1-F106L virus. (C) Single cycle growth curve with WT and VP1-F106L. HeLa cells were infected at MOI=0.1, and progeny were quantified by plaque assay. Data are mean ± SEM (n≥11, ≥3 independent experiments), *p<0.05, unpaired t-test.

### A single mutation, VP1-F106L, is sufficient for the fast replication phenotype

To uncover mutations that might confer the fast replication phenotype observed in the 4h passaged samples, we performed consensus sequencing. Viral genomes from both 4h P10 and 8h P10 samples were amplified by RT-PCR and sequenced, with a goal to identify mutations unique to the 4h P10 samples. Sequencing revealed a number of silent mutations, including several in both 4h P10 and 8h P10 samples (Table 1). Several different replicates from both the 4h P10 samples and 8h P10 samples contained the same mutations, suggesting that these mutations were likely present in our original stock and/or acquired during both fast and standard infection passages. However, a single point mutation conferring an amino acid change, phenylalanine to leucine substitution at position 106 of the VP1 capsid coding region, was unique to the 4h P10 samples (Figure 1B). The VP1-106F residue is buried in the hydrophobic core of the VP1 capsid protein.

**TABLE 1.** Consensus Sequencing reveals mutations in passaged CVB3

| **BP** | **Location** | **WT/Plasmid NT** | **NT Change** | **AA Change** | **4h P10** | **8h P10** | **Inoculum** |
| --- | --- | --- | --- | --- | --- | --- | --- |
| 1499 | VP2 | C | T | No | Yes | No | No |
| 2071 | VP3 | G | A | No | Yes | Yes | Yes |
| 2768 | VP1 | T | C (mix) | F106L | Yes | No | No |
| 3427 | 2A | G | A or G/A (mix) | No | Yes | No | No |
| 3832 | 2B | G | A | No | Yes | Yes | Yes |
| 4418 | 2C | C | C/T (mix) | No | Yes | No | No |
| 4585 | 2C | A | C | No | Yes | Yes | Yes |
| 4948 | 2C | T | C | No | Yes | Yes | Yes |
| 4951 | 2C | A | C | No | Yes | Yes | Yes |
| 5090-5091 | 3A | CG | GCA | No | Yes | Yes | Yes |
| 7271-7273 | 3'UTR | ATT | TTA | No | Yes | Yes | Yes |

To determine if the VP1-F106L mutation was sufficient for the fast-replication phenotype, we cloned this mutation into a new CVB3 infectious clone plasmid and generated VP1-F106L virus stocks. Single cycle growth curves at an MOI of 0.1 were performed using CVB3 WT P0 virus (referred to as WT) and CVB3 VP1-F106L virus. VP1-F106L replicated to higher titers than WT at 4h, 5h, 6h, and 8h post-infection (Figure 1C), indicating that this mutation was sufficient for a fast replication phenotype.

### The VP1-F106L fast replication phenotype is lost when entry is bypassed

We next determined whether the VP1-F106L mutation was sufficient for the fast-replication phenotype when viral entry was bypassed. When transfected into permissive cells, RNA from positive sense single-stranded RNA viruses is sufficient to initiate translation, RNA replication, and production of progeny viruses. Thus, transfection of viral RNA can be used to determine if a phenotype is dependent upon steps prior to viral translation, including attachment and entry. We *in vitro* transcribed RNA from WT and VP1-F106L infectious clone plasmids, transfected HeLa cells, and quantified viral progeny. Virus was collected from whole wells, including both supernatant and cell-associated virus and the titer was determined by plaque assay. Viral yields from VP1-F106L and WT were equivalent across all time points (Figure 2A). Since VP1 is capsid protein, we wondered if the VP-F106L mutation conferred altered virion assembly or egress. We performed the entry bypass experiment again, but collected culture supernatants, which reflects released virions. Again, there was no difference in viral yield across any of the time points (Figure 2B). From these experiments we conclude it is unlikely that the VP1-F106L mutant has altered viral translation, RNA replication, or assembly/egress, making it more likely that this mutation alters an early stage of the replication cycle.

**FIG 2.**
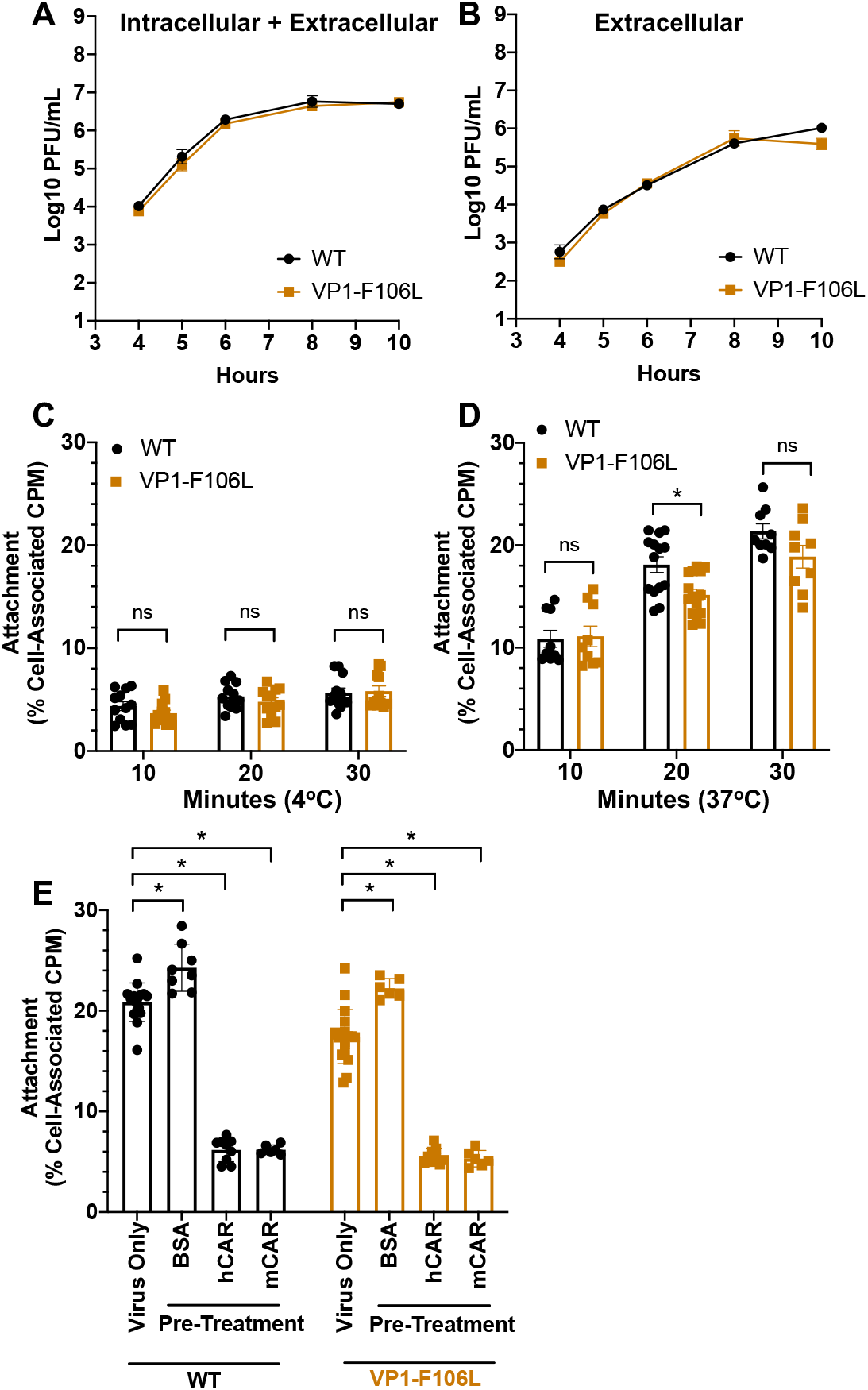
VP1-F106L has WT replication kinetics when entry is bypassed, and does not have enhanced binding to cells. (A and B) Entry bypass experiments. Viral RNA from WT and VP1-F106L was *in vitro* transcribed and transfected into HeLa cells, and progeny was quantified by plaque assay. (A) Time course for entire wells, reflecting both intracellular and extracellular progeny viruses. (B) Time course supernatants, reflecting extracellular progeny viruses. Data are mean ± SEM (n≥8, ≥3 independent experiments). Differences were not statistically significant, *p>0.05, unpaired t-test. (C-E) Viral attachment assays. Gradient purified ^35^S-labeled viruses were incubated with cells followed by washing and quantification by scintillation counting. (C) Binding assay performed at 4°C, which facilitates viral binding but not entry. (D) Binding assay performed at 37°C, which facilitates both binding and entry. (E) Binding assay performed at 37°C after pre-incubation of virus with 0.1 ug of either BSA, human CAR (hCAR), or mouse CAR (mCAR) for 15 min at room temperature. Data are mean ± SEM (n≥8, ≥3 independent experiments), *p<0.05, two-way ANOVA (multiple comparisons).

### The VP1-F106L mutation does not alter viral binding to HeLa cells

To determine whether the VP1-F106L mutation affects the first step of the replication cycle—cell attachment—we performed binding experiments with HeLa cells and ^35^S-labeled viruses. We first examined viral binding to HeLa cells at 4°C, which facilitates binding but not entry. At this temperature, both WT and VP1-F106L had equivalent binding to HeLa cells across multiple time points (Figure 2C). Next, we performed experiments 37°C, which facilitates both binding and entry. These experiments indicated that VP1-F106L did not have enhanced binding/entry and, if anything, the mutant had slightly reduced binding/entry (Figure 2D). To confirm that the ^35^S signal represents bona fide viral binding rather than background, we performed additional binding experiments where viruses were pre-incubated with BSA, human CAR (hCAR), or mouse CAR (mCAR), with the hypothesis that pre-exposure to hCAR or mCAR receptors should bind virions and inhibit ^35^S-labeled virus binding to HeLa cells. As expected, exposure to BSA did not reduce viral binding to HeLa cells, but exposure to hCAR or mCAR did (Figure 2E). These results indicate that the ^35^S signal represents bona fide viral binding to HeLa cells. Additionally, hCAR and mCAR were able to inhibit binding of both WT and VP1-F106L virus to HeLa cells. Overall, these results indicate that the enhanced replication speed of VP1-F106L is not due to enhanced attachment to HeLa cells.

### VP1-F106L virus has enhanced RNA release into HeLa cells

Given that VP1-F106L virus did not have enhanced cell attachment, we next hypothesized that the fast replication phenotype could be due to enhanced or earlier viral RNA release into cells. To test this hypothesis, we generated light-sensitive stocks of WT and VP1-F106L viruses, which can be used to monitor RNA release kinetics. Viruses were grown in the presence of neutral red dye, which is incorporated inside the viral capsid. When exposed to light, neutral red crosslinks RNA within capsids, rendering the viruses non-infectious. However, if viral RNA has been released from the capsid in the cytoplasm, the infection becomes insensitive to light inactivation due to dye diffusion away from the RNA [28]. Neutral red-labeled viruses were added to HeLa cells in the dark and plates were incubated for various amounts of time, followed by addition of an agar overlay for plaque assays. For each set of samples, one was kept in the dark to quantify total viral yield, and one was exposed to light to quantify viruses that had undergone uncoating by the time of light exposure. The ratio of light insensitive plaques to total plaques reflects the relative uncoating frequency. The VP1-F106L mutant had earlier appearance of light insensitive plaques and a greater overall frequency of light insensitive plaques over the time course compared with WT (Figure 3A). These results suggest that VP1-F106L has enhanced RNA release into HeLa cells.

**FIG 3.**
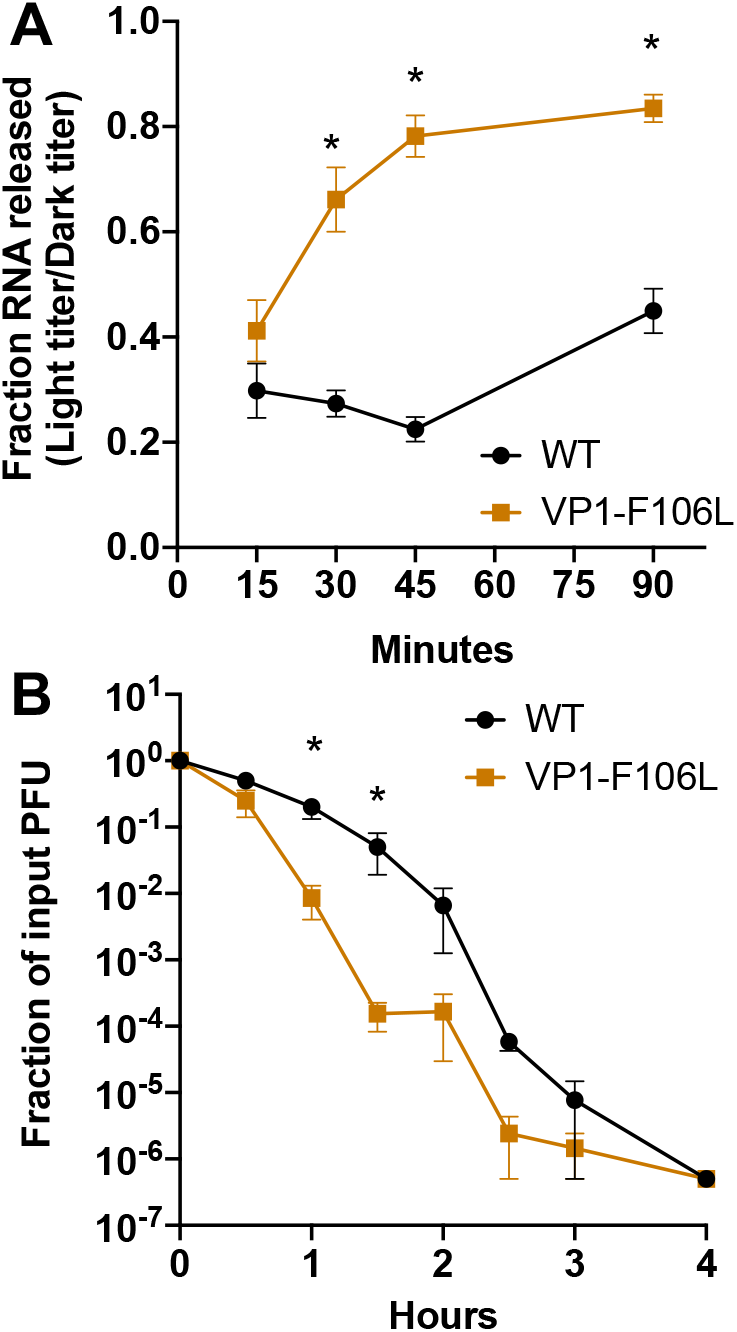
VP1-F106L has early RNA release and less stable virions. (A) Neutral-red virus RNA release assay. Virus stocks were generated with neutral red dye incorporated inside the capsid, which confers light sensitivity unless virion RNA undergoes uncoating prior to light exposure. HeLa cells were infected in the dark and incubated at 37°C, followed by washing, addition of an agar overlay, and exposure to light vs. dark at various time points. Plaques were counted and light titers were divided by dark titers to determine the ratio of virions that had released RNA by that time point. Data are mean ± SEM (n≥12, ≥5 independent experiments), *p<0.05, unpaired t-test. (B) Viral stability assay. Viruses were incubated at 44°C followed by plaque assay to quantify viable viruses remaining. Titers at 44°C were compared to control samples incubated at 4°C. Data are mean ± SEM (n≥8, ≥3 independent experiments), *p<0.05, unpaired t-test.

### VP1-F106L virions are less stable than WT

Viral RNA release requires capsid protein conformational changes and capsid instability; thus, we wondered whether VP1-F106L capsids are less stable than WT capsids. We compared WT and VP1-F106L stability at elevated temperature over time via a plaque assay. Viruses were incubated for varying amounts of time at 44°C, a temperature that reduces CVB3 capsid stability and inactivates WT particles by 4 h. These data were compared to “input” virus that was incubated at 4°C. Both WT and VP1-F106L titers were reduced by six orders of magnitude by 4 h. However, VP1-F106L virus lost infectivity significantly faster than WT at 44°C, with titers over 300-fold lower than WT at 1.5 h post-incubation (Figure 3B). These results indicate that VP1-F106L virions are less stable than WT virions.

### VP1-F106L has enhanced fitness in HeLa cells

To compare fitness of WT and VP1-F106L viruses, we performed direct competition experiments in HeLa cells. A mixture of half WT and half VP1-F106L virus was used for these experiments. HeLa cells were infected at an MOI of 0.5 using the mixed inoculum, and replication proceeded for either 4h or 8h before cells were collected. We performed 10 blind passages. Viral RNA was collected from the inoculum, passage 1 (P1), and passage 10 (P10), and the VP1 region was amplified by RT-PCR. To determine the ratio of WT to VP1-F106L, we TOPO-cloned individual RT-PCR products and performed consensus sequencing. We found that our mixed inoculum contained 42% VP1-F106L and 58% WT virus (Figure 4). However, even after just a single passage, VP1-F106L was the predominant sequence in the 4 h passage samples (62%), and, surprisingly, in the 8 h passage samples as well (77%). VP1-F106L became even more predominant with continued passaging, with 86% and 98% VP1-F016L at passage 10 for 4 h and 8 h samples, respectively (Figure 4). These data suggest that the VP1-F106L virus has fitness advantage in HeLa cells, both at early and late time points of the viral replication cycle.

**FIG 4.**
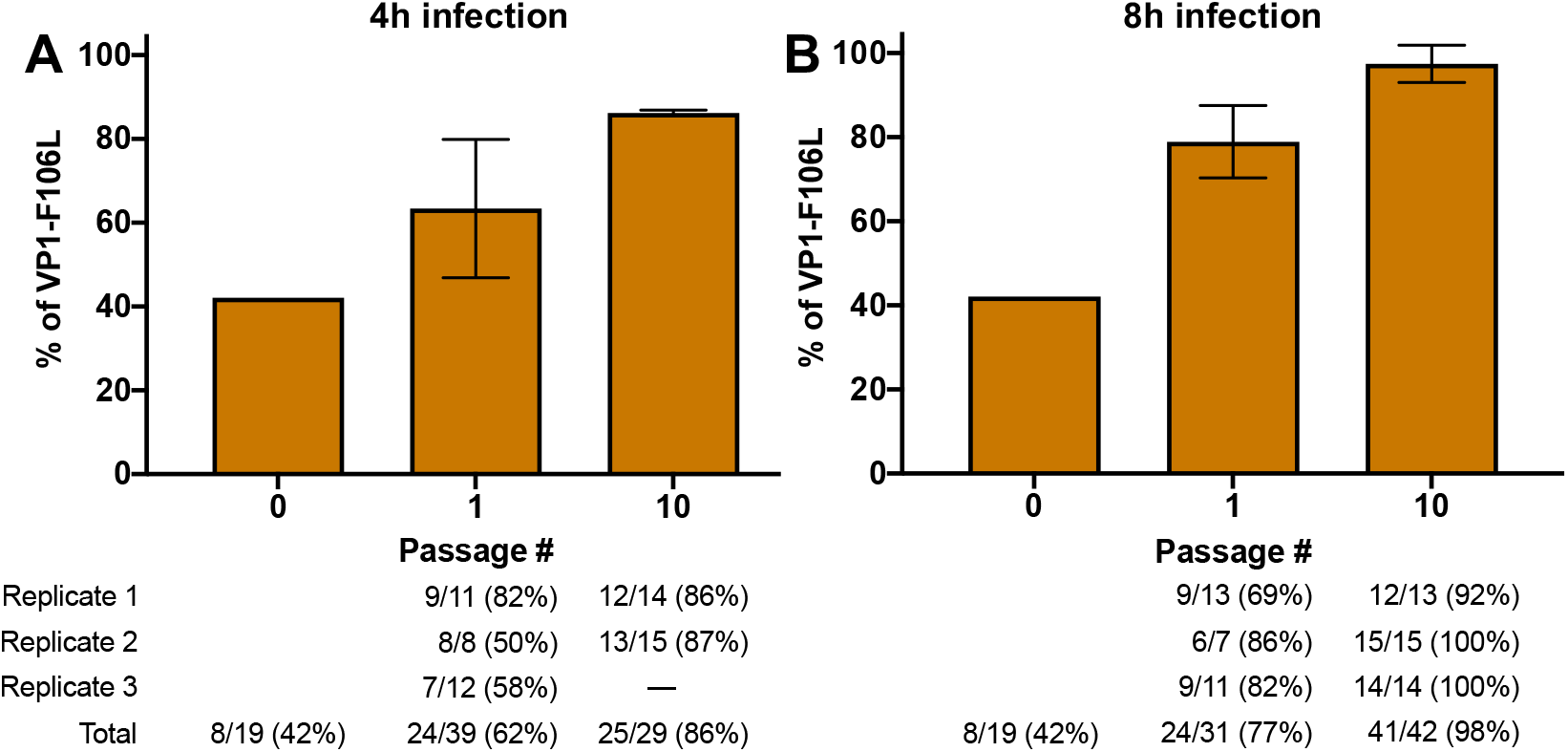
VP1-F106L is more fit that WT in HeLa cells. Direct competition experiment between WT and VP1-F106L in HeLa cells. HeLa cells were infected in triplicate with WT and VP1-F106L in a mixed inoculum at an MOI of 0.5. Progeny were harvested at (A) 4 h or (B) 8 h post infection. After 10 passages, viral RNA was collected from input, passage 1 (P1), and passage 10 (P10), and RT-PCR products were TOPO cloned and sequenced. Data are mean ± SEM (n indicated below each bar).

### VP1-F106L virus is attenuated in mice

Although VP1-F106L virus has enhanced replication speed *in vitro*, we wondered whether it would incur a fitness cost in more complex environments, such as within infected animals. To test this, we used 1 × 10^8^ PFU of WT or VP1-F106L virus to orally infect male mice lacking the interferon alpha receptor (IFNAR-/-), as these mice are more susceptible to CVB3 infection. Three days after infection, mice were sacrificed, tissues were collected, and virus was quantified by plaque assay. The VP1-F106L mutant replicated to significantly lower levels than WT in all tissues examined (Figure 5A). These differences were over 1,000-fold for several tissues, indicating that VP1-F106L virus has a strong replication defect *in vivo*. We also examined survival of mice and found that mice infected with VP1-106L virus had significantly higher survival (Figure 5B). These results indicate that VP1-106L virus is attenuated *in vivo*.

**FIG 5.**
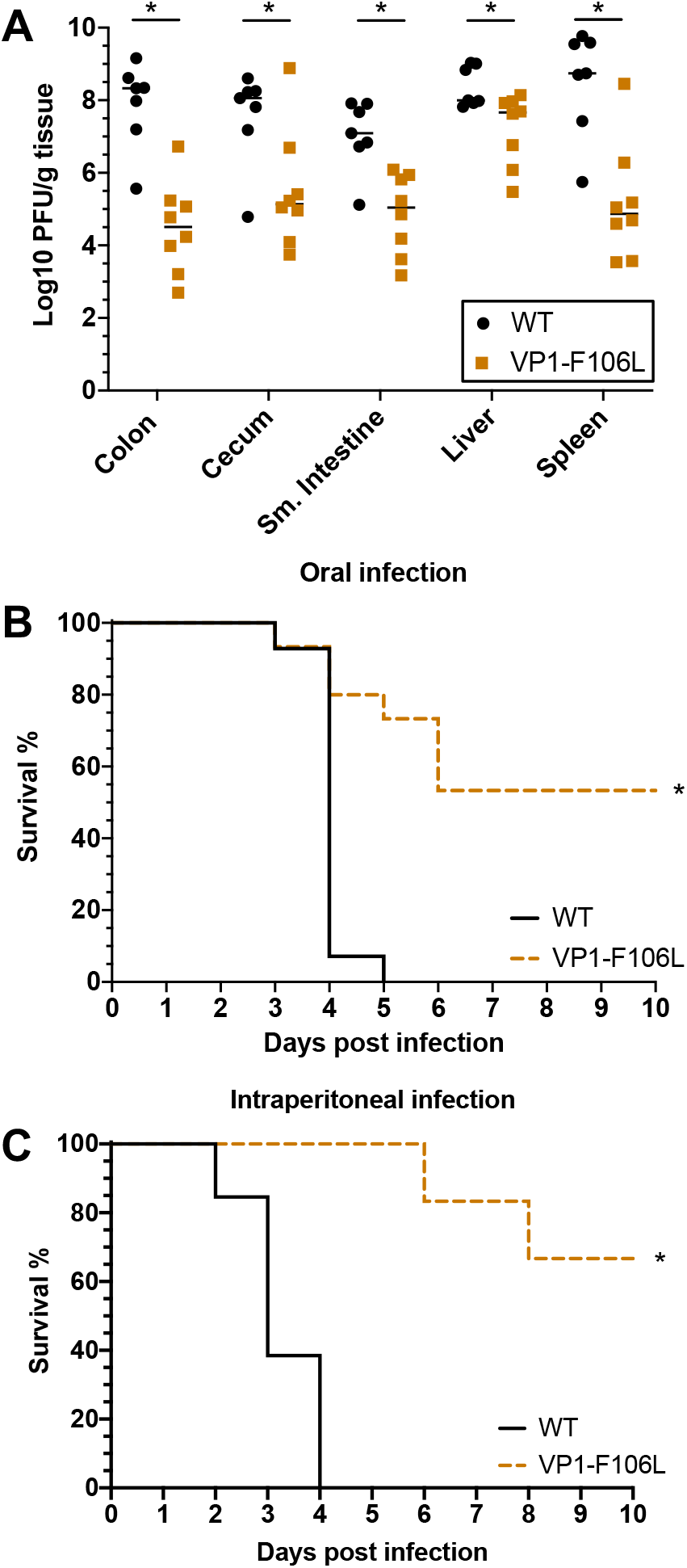
VP1-F106L is attenuated in mice. (A) Titers from orally infected IFNAR-/-mice. Male mice were orally infected with 1×10^8^ PFU of WT or VP1-F106L viruses. At three days post infection, mice were sacrificed, tissues were harvested, and titers were determined by plaque assay. Data are mean ± SEM (n≥7-8, ≥2 independent experiments), *p<0.05, unpaired t-test. (B) Survival of IFNAR-/-mice after oral infection. Mice were infected as in (A) except they were monitored twice daily for survival for 10 days. Data are represented as survival across 10 days (n≥13-14, ≥2 independent experiments), *p<0.05, log-rank test. (C) Survival of IFNAR-/-mice after intraperitoneal (IP) injection. Male mice were IP injected with 1×10^3^ PFU of WT or VP1-F106L virus. Data are represented as survival across 10 days (n≥11-13, ≥3 independent experiments), *p<0.05, log-rank test.

Due to its less stable capsid, we wondered whether VP1-F106L virus was attenuated in orally-infected mice due to the relatively extreme environments of the gastrointestinal tract, including low pH and digestive enzymes. We therefore used intraperitoneal (IP) injection to bypass the gastrointestinal lumen and compared survival of mice inoculated with 1 × 10^3^ PFU of WT vs. VP1-F106L viruses. Mice injected IP with VP1-F106L had significantly enhanced survival compared to WT, suggesting that attenuation of VP1-F106L in mice is not dependent on the oral route of infection (Figure 5C). From these studies we conclude that VP1-F106L has reduced replication and pathogenesis *in vivo*.

## Discussion

Although many viruses have been passaged in cultured cells to generate lab adapted viruses [3-7], it has been unknown whether a virus already well adapted to cell culture could evolve to replicate faster. Here, by sequential harvesting of the earliest progeny viruses, we discovered a mutant with enhanced replication speed in HeLa cells. A single mutation, VP1-F106L, was sufficient for increased replication speed. This mutation confers reduced virion stability and enhanced RNA release *in vitro*, but VP1-F106L virus is attenuated in mice. These results highlight a fitness tradeoff, where selection for enhanced replication *in vitro* leads to reduced replication *in vivo*.

Initially, we hypothesized that mutation(s) in the viral polymerase or other replication complex components would confer enhanced replication speed, but instead we found that a mutation in the VP1 capsid protein conferred the fast replication phenotype. The VP1-F016L mutation did not alter cell attachment or virion assembly, but it increased the efficiency of viral RNA release during uncoating. VP1-F106L virions were less stable than WT, suggesting that capsid instability may enhance the efficiency of RNA release during entry. These data suggest that viral RNA release is somewhat inefficient for WT virus, perhaps due to a requirement for high capsid stability *in vivo* and during transmission. While cryo-EM structures of several coxsackieviruses have revealed facets of capsid dynamics and uncoating, the efficiency of RNA release has been unclear [29-33]. Work done with the closely related poliovirus has shown that RNA release is efficient and occurs close to the cell surface at early time points after infection [34]. Our studies show that VP1-F106L CVB3 had similar kinetics of RNA release as poliovirus, but that WT CVB3 RNA release was slower [34]. In fact, WT CVB3 RNA release kinetics observed here were similar to a poliovirus mutant with increased virion stability and reduced uncoating [35]. These findings suggest that CVB3 RNA release is less efficient than poliovirus.

Viral capsids must balance the need for stability in a variety of environments to ensure transmission, while maintaining the ability to fall apart upon receptor engagement and uncoating [35-38]. Further studies will be required to investigate the exact mechanism by which WT and VP1-F106L differ in capsid stability and RNA release. There are conflicting reports about role of endosomal acidification in CVB3 entry into HeLa cells [24, 39]. It is possible that WT and VP1-F106L have different pH sensitivities that alter RNA release. During CAR binding, the CVB3 particle undergoes a structural change to prime RNA release [40, 41]. The F106 residue of VP1 is in a hydrophobic pocket near the site of RNA release, so it is plausible that VP1-F106L virions more closely resemble the CVB3 alpha-particle and is primed for RNA release.

While VP1-F106L had increased fitness *in vitro*, it had reduced fitness and virulence in mice. The attenuation of VP1-F106L in mice could be due to several different factors. It is likely that VP1-F106L is attenuated due to reduced viral capsid stability, a finding that is well characterized across many different viral families [37, 42, 43]. The reduced fitness of VP1-F106L in orally infected mice is not simply due to virion instability in the low pH of the stomach, since the mutant was also attenuated in mice injected intraperitoneally, bypassing the gut lumen. It is also possible that VP1-F106L is more sensitive to immune responses in mice. More efficient RNA release into cells could be disadvantageous to the virus in an animal model. VP1-F106L early release of free RNA into the cytoplasm could be sensed to activate innate immune responses [44]. The production of Type-II IFN by other RNA sensing proteins such as TLR3 has been shown to be crucial for protection against CVB3 [45, 46]. Another possibility is that VP1-F106L releases RNA into the endosome that can be sensed by TLR7 or TLR8 to lead to macrophage or dendritic cell activation [47-49]. These innate RNA sensing mechanisms likely play a larger role *in vivo* than *in vitro*, particularly due to the longer time frame of the mouse infections.

This work is one example of how a single mutation can lead to fitness tradeoffs for an enteric virus. We show that a cell-adapted virus strain can be passaged to generate a mutant that replicates faster than WT virus but has reduced fitness in mice. It highlights that different selective pressures shape viral populations in different environments.

## Methods

### Cells and virus stocks

HeLa cells were maintained in DMEM media with 10% calf serum. CVB3-H3 (from Marco Vignuzzi) was generated by transfecting HeLa cells with an infectious clone plasmid and a plasmid expressing T7 RNA polymerase, followed by two rounds of amplification to generate high titer stocks. Virus titers were determined by plaque assay using HeLa cells [50].

### Virus passaging experiments

HeLa cells were plated at a density of 3 × 10^6^ cells/well on 6 well dishes and cells were infected using a MOI of 0.5. Virus was absorbed for 30 minutes at 37°C, the supernatant was aspirated, cells were washed twice, and 1mL of media was added. Infection was allowed to continue for 4 hours or 8 hours at which point cells were scraped into the media and the entire 1 mL sample was collected. Samples were freeze/thawed three times to release any intracellular virus, and 500 ul of the sample was used to begin a new infection. Viral titers were determined by plaque assay after passage 5, and passage 6 was initiated with a MOI of 0.5, followed by several additional passages of unknown MOI as above. These gain-of-function BSL2 virus selection experiments were approved by the UT Southwestern Institutional Biosafety Committee.

### Sequencing

Virus collected from passage 10 was used to infect HeLa cells, and viral RNA was isolated with TriReagent and amplified by RT-PCR as previously described [50]. PCR products were purified with the Nucleospin Gel and PCR Clean-up kit (Macherey-Nagel). Sequencing was performed by the McDermott Sequencing Center at UT Southwestern with the primers listed in Supp. Table 1. RT-PCR of the VP1 region was performed using primer 15R for cDNA synthesis, and primers 7F and 7bR for PCR, and primers 6R and 7F for sequencing (see Supp. Table 1 and Figure 4)

### Single cycle growth curves

1 × 10^5^ HeLa cells/well in 96-well plates were infected with 1 × 10^4^ PFU/well for 30 min, the supernatant was aspirated, cells were washed twice, and 100 uL of media was added. At each timepoint plates were moved to -80°C, followed by freeze/thawing three times to release any intracellular virus, and viral titers were determined by plaque assay. These titers reflect combined extracellular and intracellular virus at each timepoint.

### Cloning of VP1-F106L mutant

KLD site directed mutagenesis was used to create a pCVB3-H3 plasmid with the VP1-F106L mutation. PCR was performed using Q5 polymerase (New England Bioloabs) and the following primers: (sense) 5’-AAAGCTAGAAcTCTTTACCTACG-3’ and (antisense) 5’-CTCCTAAGTTGTGCCGCT-3’. The product was digested using the KLD Enzyme Mix reagents (New England Biolabs) and transformed into DH5-alpha competent cells. The entire CVB3 region of the plasmid was sequenced to confirm the presence of the VP1-F106L mutation and the absence of any other mutations. VP1-F106L virus was generated from this plasmid as described above.

### Entry bypass assay

pCVB3-H3-WT and pCVB3-H3-VP1-F106L plasmids were linearized with ClaI and purified with the QIAquick Gel Extraction kit (Qiagen). *In vitro* transcription was performed with the P1230 kit (Promega). 2ug of viral RNA was transfected into 1 × 10^6^ cells HeLa cells using Lipofectamine 2000 (ThermoFisher). At each time point, the cells were collected by scraping, and samples were freeze-thawed three times to release intracellular virus. Viral titers were determined by plaque assay.

### Viral stability assays

1 × 10^5^ PFU of virus was added to 300ul PBS+ (PBS supplemented with 100ug/ml CaCl_2_and 100ug/ml MgCl_2_) in a glass tube followed by incubation at 4°C (to generate input values) or 44°C. Viral titers after incubation were determined by plaque assay using HeLa cells.

### Neutral red virus production and RNA release assays

To create light-sensitive viruses for the RNA release assay, WT and VP1-F106L viruses were grown in the dark in HeLa cells supplemented with 50 ug/mL neutral red dye [51]. Cells were freeze/thawed three times and viral titers were determined by plaque assays performed both in the dark and in light. The amount of light-insensitive background titer was negligible: Less than 1 in 1.5×10^5^ PFU of each stock was light-insensitive.

For RNA release assays, HeLa cells were plated at a density of 3 × 10^6^ cells/well. In the dark, 500 PFU per well was added and plates were incubated at 37°C. At the indicated time points, virus was aspirated, cells were washed twice, and an agar overlay was added to all samples in the dark. Immediately after adding the overlay, paired samples were exposed to light or dark at room temperature for 10 minutes. After 10 minutes, all plates were incubated at 37°C until plaques were counted two days later.

### ^35^S-labeled virus production and binding assays

^35^S-labeled virus stocks for binding assays were generated as previously described [52]. Briefly, cells were pulsed with ^35^S-Cysteine/Methionine during infection to generate radioactive virus stocks, which were then purified by CsCl gradient ultracentrifugation. Purity was confirmed by running the samples on gels followed by phosphorimaging.

Binding assays were performed as previously described [50]. Briefly, HeLa cells in 6-well dishes were incubated with 5000 CPM ^35^S-labeled virus at 4°C or 37°C, followed by aspiration of the virus, washing twice, and cells were collected. For samples that were mixed with protein (BSA, human CAR, mouse CAR [Fisher Scientific]) prior to attachment, virus was incubated with 0.1ug of protein for 15 min at room temp before addition to HeLa cells. Input samples were incubated at 4°C for the duration of the experiment and scintillation counted at the end of the experiment with other samples.

### Animal experiments

All animals were handled according to the Guide for the Care of Laboratory Animals of the National Institutes of Health. All mouse studies were performed at UT Southwestern (Animal Welfare Assurance no. a3472-01) by using protocols approved by the local Institutional Animal Care and Use Committee in a manner designed to minimize pain, and any animals that exhibited severe disease were euthanized immediately with isoflurane. C57BL/6 mice defective for the interferon alpha/beta receptor (IFNAR-/-) were obtained from Jackson Laboratories and all experimental mice were 8-to 12-weeks old. Due to sex bias in CVB3 infection, only male mice were used in experiments [53]. Mice were perorally infected with 1 × 10^8^ PFU of virus. Feces were collected for three days after infection, at which time mice were sacrificed and tissues were collected. Tissues were homogenized by bead beating with PBS (Next Advance Bullet Blender Storm) and titers were determined by plaque assay. In separate experiments, mice were perorally infected with 1 × 10^8^ PFU of virus and survival was monitored. For some experiments, mice were intraperitoneally injected with 1 × 10^3^ PFU of virus and survival was monitored.

### Competition experiments

HeLa cells were infected at an MOI of 0.5 with 50% WT CVB3 and 50% VP1-F106L CVB3 in triplicate. As controls, wells were also infected with 100% WT or 100% VP1-F106L in triplicate. Virus was allowed to replicate for either 4h or 8h and the entire well (1mL) was collected by scraping. Samples were freeze-thawed three times and cellular debris was removed by centrifugation. Supernatant (500ul) was used to infect new cells. This process was repeated 10 times. After 10 passages, samples were used to infect HeLa cells, and RNA was collected at 8h post-infection using TriReagent. The VP1 region was amplified by RT-PCR and PCR products were cloned using TOPO-TA Cloning Kits (ThermoFisher) (Primers used: 15R, 7F, 7bR, and 6R listed in Supp. Table 1). Plasmids from individual colonies were sequenced as described above to determine the ratio of WT to VP1-F106L.

## Acknowledgements

We thank Melissa Budicini and Andrea Erickson for critical review of the manuscript, Wenchun Fan for advice on entry bypass experiments, and Robert Maples for training on animal experiments. Work in J.K.P.’s lab is funded through NIH NIAID grant R01 AI74668, a Burroughs Wellcome Fund Investigators in the Pathogenesis of Infectious Diseases Award, and a Faculty Scholar grant from the Howard Hughes Medical Institute. ML was supported in part by NIH GM109776 Molecular Medicine NRSA Institutional Predoctoral T32 Training Grant.

---

**Supplemental Table 1.**
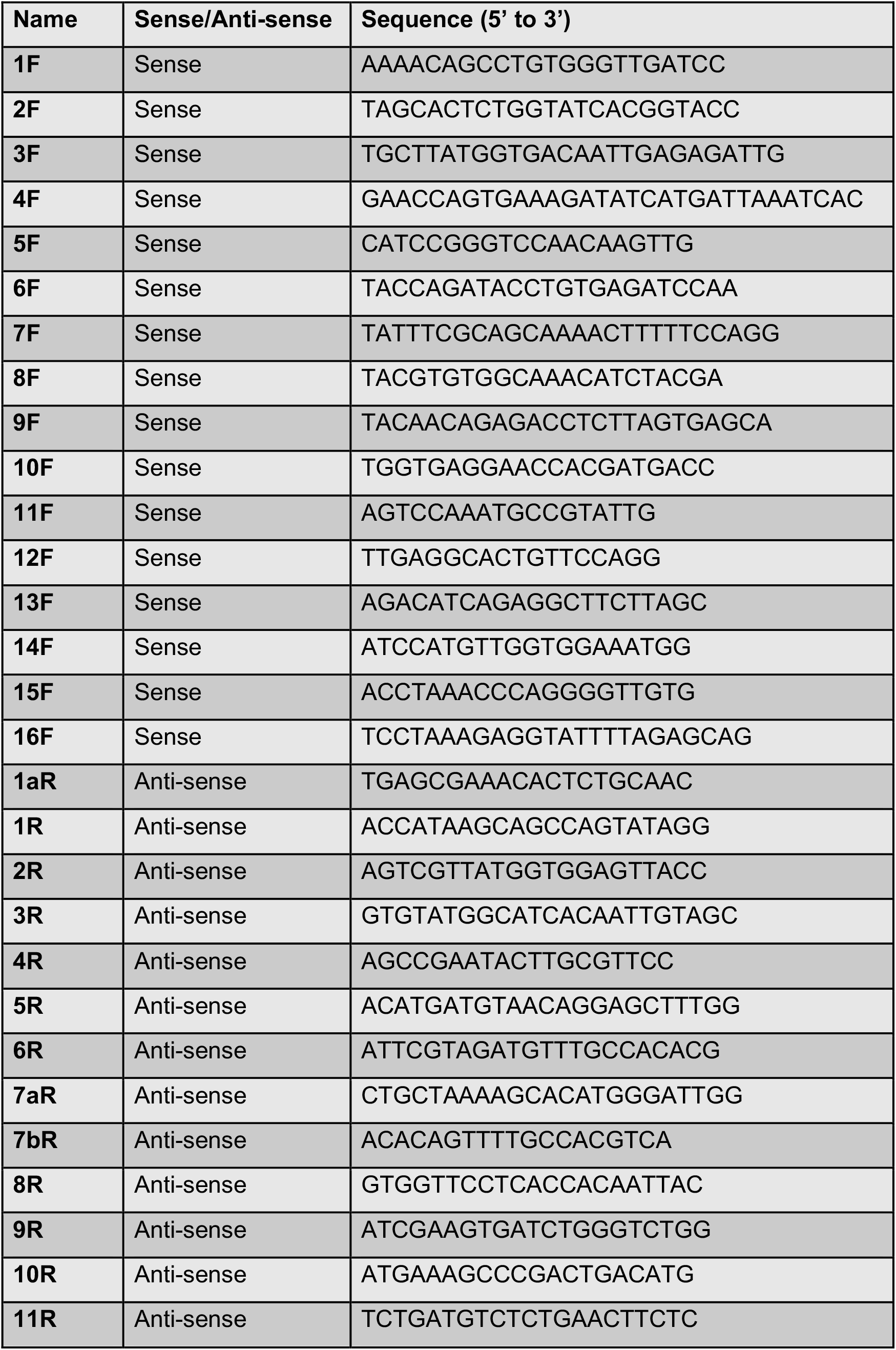
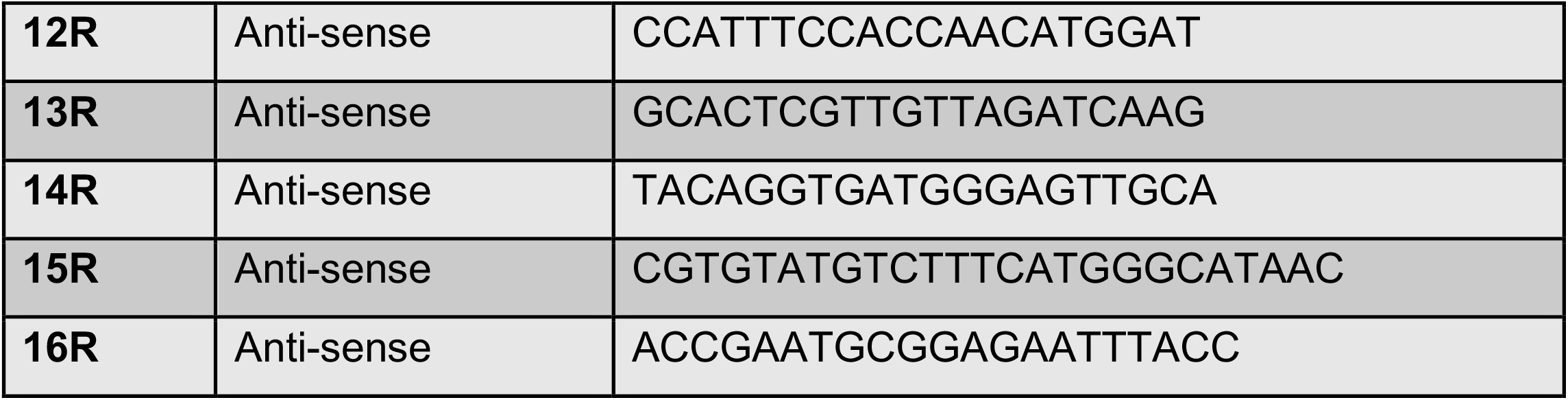
Primers used for sequencing

